# A high resolution 7-Tesla resting-state fMRI test-retest dataset with cognitive and physiological measures

**DOI:** 10.1101/008706

**Authors:** Krzysztof J. Gorgolewski, Natacha Mendes, Domenica Wilfling, Elisabeth Wladimirow, Claudine J. Gauthier, Tyler Bonnen, Florence J. M. Ruby, Robert Trampel, Pierre-Louis Bazin, Roberto Cozatl, Jonathan Smallwood, Daniel S. Margulies

**Affiliations:** Max Planck Research Group for Neuroanatomy and Connectivity, Max Planck Institute for Human Cognitive and Brain Sciences, Leipzig, Germany; Department of Neurophysics, Max Planck Institute for Human Cognitive and Brain Sciences, Leipzig, Germany; Department of Neurology, Max Planck Institute for Human Cognitive and Brain Sciences, Leipzig, Germany; Concordia University/PERFORM Center, Montreal, Canada; Department of Psychology, University of York, York, United Kingdom; Databases and IT Group, Max Planck Institute for Human Cognitive and Brain Sciences, Leipzig, Germany

## Abstract

Here we present a test-retest dataset of functional magnetic resonance imaging (fMRI) data acquired at rest. 22 participants were scanned during two sessions spaced one week apart. Each session includes two 1.5 mm isotropic whole-brain scans and one 0.75 mm isotropic scan of the prefrontal cortex, giving a total of six timepoints. Additionally, the dataset includes measures of mood, sustained attention, blood pressure, respiration, pulse, and the content of self-generated thoughts (mind wandering). This data enables the investigation of sources of both intra- and inter-session variability not only limited to physiological changes, but also including alterations in cognitive and affective states, at high spatial resolution. The dataset is accompanied by a detailed experimental protocol and source code of all stimuli used.

## Background and Summary

In contrast to the focused scope of task-based functional studies using functional magnetic resonance imaging (fMRI), data acquired independent of specific task demands has become the basis for post hoc investigation of diverse functional systems. Rather than constraining the question of functional organization to a single paradigm, the observation that ongoing intrinsic activity replicates networks observed during task has made resting-state fMRI (rs-fMRI) data a highly versatile resource.

To date the vast majority of resting-state fMRI research has been conducted using data acquired with a resolution of approximately 3-4 mm^3^. This voxel size provides a judicious compromise between signal-to-noise ratio and spatial resolution when using a standard 3-Tesla MRI scanner. However, as the thickness of gray matter in the cerebral cortex can range from 1.5 to 3 mm, partial sampling of tissue becomes a substantial source of noise when using the standard voxel size. Although traditionally attenuated using techniques such as spatial smoothing, such approaches are not only considered increasing problematic for investigating brain function, but also reduce the resolution with which neuroanatomical questions can be investigated.

7 Tesla (7T) ultra-high field MRI scanners, though not as common as their 3 Tesla (3T) counterparts, offer notably enhanced spatial resolution and are becoming increasingly incorporated into academic and clinical research. The major advantage of using a field strength of 7T scanner is the resulting high signal-to-noise ratio (SNR) which can be directly translated into a high spatial resolution. Initial studies have shown that using 7T allows for the more precise delineation of resting state networks than with 3T, especially if using small voxels with edge lengths in the range between 1 and 1.5 mm ^1–3^. It was further shown that spontaneous neuronal activity is one of the major contributors to the measured fMRI signal fluctuations at 7T, increasing almost twofold relative to earlier experiments under similar conditions at 3T ^4^. Hence, using the increased BOLD sensitivity of ultra-high fields for resting state-fMRI, a number of studies with different neuro-scientific questions were conducted at 7T ^5,6^. However, also an increased propensity for artifacts at 7T compared to 3T was reported for resting state fMRI ^7,8^. It is therefore essential to investigate the reliability of rs-fMRI at 7T. Thus, in this article, we present the first extensive 7T test-retest dataset in order to address this issue.

A growing body of research continues to establish the reliability of various analytic approaches to rs-fMRI data^9–12^. While these studies have established rs-fMRI as a viable method for scientific inquiry, there are still methodological questions to be answered. To understand reliability within the context of rs-fMRI paradigms, it is important not only to assess the between session variance, but also to attribute that variance to distinct confounding factors^13,14^. More specifically, these factors might be hardware-specific, physiological, or mental state-related in nature. It has been shown, for example, that participant motion increases between session variance^13,15^. The following dataset allows researchers to address these questions, by including information about participant motion (derived from imaging data), breathing, pulse, and blood pressure. Additionally, to assess the influence of changing the position of the head of the participant within the coil, the participants were taken out, and then returned to the scanner within the same scanning session.

This dynamic aspect of spontaneous brain fluctuations has recently been the focus of intense methodological development^16–18^. Differences in mental state, mood, and content of spontaneously generated thoughts during the scanning session may translate into increased between-session variance that is independent of physiological factors. To address these uncertainties, the dataset includes assessments of the participants’ mood and sustained attention immediately prior to each scanning session. The content of self-generated thoughts immediately after each resting-state run was also probed using an in-scanner adaptation of the *New* York Cognition Questionnaire^19^. This data allows researchers to relate dynamic changes in brain states to externally measured differences in mental states.

Additionally, reliability of an analysis is not only restricted to experimental properties of the data. The MRI field has developed a variety of data processing improvements focusing on data cleaning (removal of confounds), normalization (transforming brains of each individual into a common space), and feature selection (deciding which aspect of spontaneous activity the analysis should focus on). Those processing steps can have a significant influence on the ultimate reliability of the results. Access to high resolution and quality test-retest data will allow authors of tools and methods to test their performance in terms of reliability. High resolution data is especially important for methods dealing with interindividual differences and investigating cortical layer-dependent processes. Additionally reliability metrics such as the Intra-class Correlation Coefficient^20^ make the assumption about the spatial correspondence of each voxel between subjects. High resolution datasets like this one can reveal to what extent this assumption is just an approximation.

We present a dataset consisting of high resolution brain scans — including 0.7 mm^3^ MP2RAGE anatomical scans — collected from 22 individuals at two time points. This dataset is optimal for addressing methodological questions requiring reliability assessment, and also enables the investigation of different contributing confounds to between-session variance. In addition to the crucial need for methods validation specific to high resolution data, the relation of mental state (using measures of mood, sustained attention, and the content of self-generated thought) to intrinsic brain dynamics is a foundational question of neuroscience for which we hope this dataset will provide insight.

## Methods

### Participants

22 participants (10 women) were selected from a database of people having previously taken part in 7T experiments at the Max Planck Institute for Human Brain and Cognitive Sciences, Leipzig, Germany. All participants were native German speakers. Their age ranged from 21 to 29 with mean 25.1. All of the participants have previously taken part in MRI experiments at the 7T facility (from 5 to 51 times, mean 23) and were therefore accustomed with the procedure. All subjects had given written informed consent and the study was approved by the Ethics Committee of the University of Leipzig.

### Testing procedure

Each participant was invited to the institute twice exactly one week apart (therefore visits were matched in terms of time of the day and day of the week). A similar testing protocol was applied to the participants on both visits. Participants were instructed to refrain from drinking caffeinated products starting two hours before each visit. They were reminded of this requirement via a text message sent on the day of the experiment. The data acquired is summarized in Figure 1, and the order of tasks/measurements was the following (details of each test are described in the next section):

**First visit:**

1. At the behavioural testing room:
  a. Briefing and consent signature
  b. Conjunctive Continuous Performance Task (CCPT)
  c. Mini *New* York Cognition Questionnaire (mini NYC-Q)
  d. Positive and Negative Affect Schedule - Expanded Form Questionnaire (PANAS-X)
  e. Hydration, caffeine, sleep questionnaire
  f. 5 minutes of rest
  g. Blood pressure and pulse measurements
2. At the scanner
  a. Instructions for the in-scanner version of CCPT
  b. Localizer scan
  c. Structural scan
  d. Field map scan
  e. Whole-brain resting state scan (15min)
  f. In-scanner mini NYC-Q
  g. Field map scan
  h. Whole-brain resting state scan (15min)
  i. In-scanner mini NYC-Q
  j. Prefrontal cortex submillimeter scan (10min)
  k. In scanner mini NYC-Q

Second visit:

1. At the behavioural testing room - same as the first visit
2. At the scanner - differences from the first visit in bold:
  a. Instructions for the in scanner version of CCPT
  b. Localizer scan
  c. Field map scan
  d. Whole-brain resting state scan (15min)
  e. In scanner mini NYC-Q
  f. **Take participant out of the scanner, ask them to sit in an upright position and put them back into the scanner**
  g. **Localizer scan**
  h. Field map scan
  i. Whole-brain resting state scan (15min)
  j. In-scanner mini NYC-Q
  k. Prefrontal cortex submillimeter scan (10min)
  l. In-scanner mini NYC-Q

**Figure 1.**
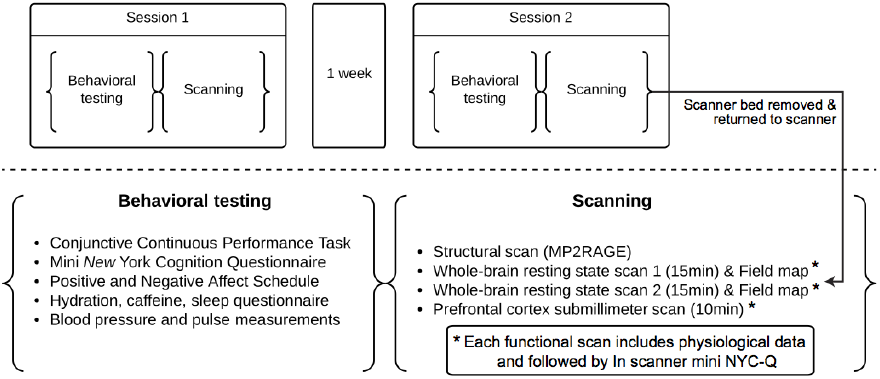
**Summary of the acquired data for two sessions of behavior testing and MRI scanning.**

### Behavioural tests

#### Conjunctive Continuous Performance Task

The Conjunctive Continuous Performance Task looks at the ability to sustain attention over a period of approximately 20 min.^21^. The task involves looking at series of shapes (triangles, squares, circles, and stars) in different colors (red, green, blue, yellow). The participant is asked to react by pressing the spacebar of a computer keyboard if and only if they see a red square (which appears 30% of the time). Each shape is displayed for 100ms and is separated from the next by a randomly-selected inter-stimulus interval (ISI) of 1000, 1500, 2000, or 2500. There was no feedback. The test consisted of 320 trials preceded by 15 trials of a practice run (which did include immediate feedback). After the practice run and before the test proper the examiner made sure that each participant understood the task. All the instructions on the screen were presented in German. The task was implemented in OpenSesame^22^ using PsychoPy backend^23,24^ - the code is available at https://github.com/NeuroanatomyAndConnectivity/ConjunctiveContinuousPerformanceTask For more details of the task please see the original paper^21^.

#### Mini New York Cognition Questionnaire

The mini version of the *New* York Cognition Questionnaire is an adaptation of the full version of *New* York Cognition Questionnaire^19^. The aim of the mini version was to shorten the questionnaire based on factors extracted from the full version (see ^19^ for details). The questionnaire consists of series of statements, each starting with “During the last measurement”:

1. I thought about something positive
2. I thought about something negative
3. my thoughts involved future events
4. my thoughts involved past events
5. my thoughts involved myself
6. my thoughts involved other people
7. my thoughts involved my surroundings
8. I was fully awake
9. my thoughts were in the form of images
10. my thoughts were in the form of words
11. my thoughts were more specific than vague
12. my thoughts were intrusive

Participants had to rate each statement on a visual analog scale ranging from “Completely did not describe my experience” (score 0, left hand side) to “Completely described my experience” (score 100, right hand side). All questions and instructions were presented in German. The questionnaire was implemented in OpenSesame^22^ using PsychoPy backend^23,24^ as a slider operated with three buttons (left, right, next question). The slider range was from 0 to 100 with increments of 5. The initial position of the slider was 50. Question order was randomized at each presentation. The questionnaire was presented immediately after the CCPT task, and after each resting state scan (a special version of the task operated with a response box was used inside the scanner). The code of the questionnaire is available at https://github.com/NeuroanatomyAndConnectivity/NYC-Q.

#### Positive and Negative Affect Schedule - Expanded Form Questionnaire

Positive and Negative Affect Schedule - Expanded Form Questionnaire is a questionnaire probing the mood of participants^25^. It consists of 60 items which are broken down into several subscales such as General Positive and General Negative Affect. For each item (an adjective such as “happy” or “alone”) participants had to rate “to what extent have they felt this way during the past week.” This was on a scale from “very slightly or not at all” (1) to “extremely” (9). We have used a German version of the questionnaire first introduced by Gruhn et al.^26^. The questionnaire was presented using the LimeSurvey software (http://www.limesurvey.org/).

#### Hydration, caffeine, sleep questionnaire

To assess sleep patterns, hydration, and caffeine intake, participants were asked to reply to a series of additional questions.

1. On average, how many hours do you sleep every night?
2. How many hours did you sleep last night?
3. How well rested do you feel right now? 1 (Extremely tired) - 9 (Perfectly well rested)
4. How well did you sleep last night? 1 (I slept terribly) - 9 (I slept very well)
5. How well hydrated do you feel right now? 1 (Completely dehydrated) - 9 (Perfectly hydrated)
6. On average, how much water (and other liquids) do you drink every day (in litres)?
7. Comparing to other days did you drink more or less water today? 1 (I drank much less than usual) - 5 (I drank the same amount as usual) - 9 (I drank much more than usual)
8. On average, how much caffeinated drinks (coffee, cola, club mate etc.) do you drink every day (in litres - one cup = 0.2 l)?
9. Comparing to other days did you drink more or less coffee and other caffeinated drinks (coffee, cola, club mate etc.) today? 1 (I drank much less than usual) - 5 (I drank the same amount as usual) - 9 (I drank much more than usual) This questionnaire was also presented using LimeSurvey software (http://www.limesurvey.org/).

#### Blood pressure and pulse measurement

After filling the questionnaires participants were asked to relax for five min. in order for their heart rate to stabilize. After this period, their systolic and diastolic blood pressure, as well as pulse, were measured for both arms. The measurement was performed using Omron M500 device (Omron Healthcare Co. Ltd.).

### Magnetic Resonance Imaging

All experiments were performed on a 7T whole-body MR scanner (MAGNETOM 7T, Siemens Healthcare, Erlangen, Germany). A combined birdcage transmit and 24 channel phased array receive coil (NOVA Medical Inc, Wilmington MA, USA) was used for imaging. During the scan the participants’ pulse was monitored using a pulse oximeter. Breathing was measured using a pneumatic sensor. Both breathing and pulse signals were recorded using Biopac MP150 system (Biopac Systems Inc., USA) sampled at 5000 Hz with the Biopac Acqknowledge 4.1 software (Biopac Systems Inc., USA). Participants filled the mini NYC-Q questionnaire using a four button response box held in both hands. The position of the slider was adjusted by the left and right buttons, pressing any of the two middle buttons advanced to the next question. Acquisition parameters of the relevant sequences are summarised below.

#### Structural scan

For structural images a 3D MP2RAGE^27^ sequence was used: 3D-acquisition with field of view 224 x 224 x 168 mm^3^ (H-F; A-P; R-L), imaging matrix 320 x 320 x 240, 0.7 mm^3^ isotropic voxel size, Time of Repetition (TR) = 5.0 s, Time of Echo (TE) = 2.45 ms, Time of Inversion (TI) 1/2 = 0.9 s / 2.75 s, Flip Angle (FA) 1/2 = 5° / 3°, Bandwidth (BW) = 250 Hz/Px, Partial Fourier 6/8, and GRAPPA acceleration with iPAT factor of 2 (24 reference lines).

#### Field map

For estimating B0 inhomogeneities, a 2D gradient echo sequence was used. It was acquired in axial orientation with field of view 192 x 192 mm^2^ (R-L; A-P), imaging matrix 64 x 64, 35 slices with 3.0 mm thickness, 3.0 mm^3^ isotropic voxel size, TR = 1.5 s, TE1/2 = 6.00 ms / 7.02 ms (which gives delta TE = 1.02 ms), FA = 72°, and BW = 256 Hz/Px.

#### Whole-brain rs-fMRI

Whole-brain resting state scans were acquired using a 2D sequence. It used axial orientation, field of view 192 x 192 mm^2^ (R-L; A-P), imaging matrix 128 x 128, 70 slices with 1.5 mm thickness, 1.5 mm^3^ isotropic voxel size, TR = 3.0 s, TE = 17 ms, FA = 70°, BW = 1116 Hz/Px, Partial Fourier 6/8, GRAPPA acceleration with iPAT factor of 3 (36 reference lines), and 300 repetitions resulting in 15 minutes of scanning time. Before the scan subjects were instructed to stay awake, keep their eyes open and focus on a cross. In order to avoid a pronounced g-factor penalty ^28^ when using a 24 channel receive coil, the acceleration factor was kept at a maximum of 3, preventing the acquisition of whole-brain data sets at submillimeter resolution. However, as 7T provides the necessary SNR for such high spatial resolutions a second experiment was performed with only partial brain coverage but with an 0.75 mm isotropic resolution.

#### Prefrontal cortex rs-fMRI

The submillimeter resting state scan was acquired with a zoomed EPI^29^ 2D acquisition sequence. It was acquired in axial orientation with skewed saturation pulse^30^ suppressing signal from posterior part of the brain (see Figure 2). The position of the field of view was motivated by the involvement of medial prefrontal cortex in the default mode network and mindwandering^31^. This location can also improve our understanding of functional anatomy of the prefrontal cortex which is understudied in comparison to primary sensory cortices. Field of view was 150 x 45 mm^2^ (R-L; A-P), imaging matrix = 200 x 60, 40 slices with 0.75 mm thickness, 0.75 mm^3^ isotropic voxel size, TR = 4.0 s, TE = 26 ms, FA = 70°, BW = 1042 Hz/Px, Partial Fourier 6/8. A total of 150 repetitions were acquired resulting in 10 minutes of scanning time. Before the scan subjects were instructed to stay awake, keep their eyes open and focus on a cross.

**Figure 2.**
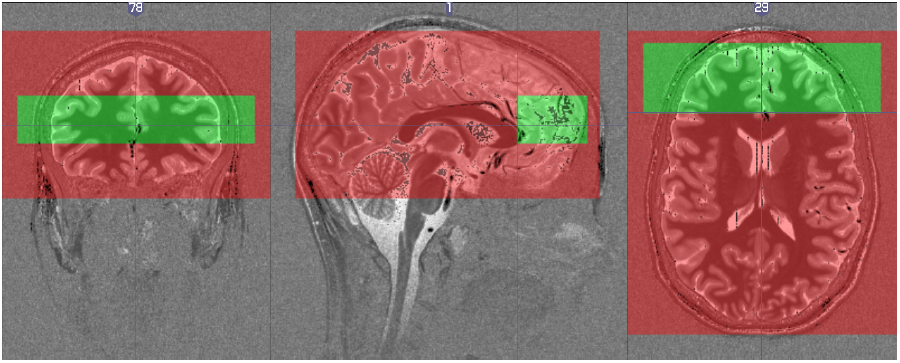
Field of view of the whole-brain resting state scan (red) and prefrontal cortex resting state scan (green). Fields of view of the two scans were overlaid on top of a T1 map for one participant. Note that whole-brain scan does not fully cover the cerebellum and brainstem.

It has to be mentioned that for both resting state experiments only a single slice was excited at a time. Although simultaneous-multi-slice imaging enables a significant reduction in repetition time, a certain coil geometry is necessary for its successful application^32^. Preliminary tests with our 24 channel coil showed that the somewhat limited number of receive elements in head-feet direction prevented an artifact-free implementation of multi-band acquisition techniques as it would be possible with, e.g., a 32 channel coil. The lack of multislice acquisition resulted in relatively slow repetition time (3 and 4 s for full brain and prefrontal cortex respectively). This unfortunately limits the use of this dataset to investigating brain dynamics (and their physiological correlates) at lower frequencies.

## Data Records

All data records listed in this section are available from the INDI/CoRR consortium facilitated through the COINS database^33^ (DOI:10.15387/fcp_indi.corr.mpg1). A README file with a detailed description of the content of all downloads is also available at this URL. The data is also available at http://openscience.cbs.mpg.de/7t_trt as a standard FTP download as well as BitTorrent facilitated by Academic Torrents to improve distribution^34^. Additionally the data has been deposited in our institute’s XNAT^35^ instance available publicly at http://xnat.cbs.mpg.de. We strongly encourage all users to subscribe to the http://groups.google.com/group/7t_trt mailing list for future updates and announcements.

All DICOM files were anonymized to remove any information that could identify the participants. The data set includes both the DICOM files as well as NIFTI files. DICOM files were converted to the NIfTI format using the dcmstack converter (https://github.com/moloney/dcmstack). All DICOM metadata were saved in the header of the NIFTI files in JSON format.

### Demographics

Location: demographics.csv

File format: plain text, comma separated values

Basic demographic information (sex, age at the first scan in years, handedness, and number of 7T scans previously taken) is available in a comma-separated value (CSV) file.

### Questionnaires and blood pressure measurements

Location: questionnaires_and_blood_pressure.csv

File format: plain text, comma separated values

Participants’ responses to the PANAS-X, sleep, caffeine, hydration questionnaires as well as the pre scan blood pressure are available in a CSV file. Data are structured as two lines per participant (one line per visit per participant) with questionnaire items as columns. A description of all items is given in Table 1.

**Table 1.** Description of the columns in the questionnaires_and_blood_pressure.csv file.

| **Column name** | **Description** | **German translation (if applicable)** |
| --- | --- | --- |
| subject_id | Participant ID | n/a |
| session | Which visit the score corresponds to | n/a |
| panas_cheerful | PANAS-X item | fröhlich |
| panas_disgusted | PANAS-X item | angeekelt |
| panas_attentive | PANAS-X item | aufmerksam |
| panas_bashful | PANAS-X item | scheu |
| panas_sluggish | PANAS-X item | träge |
| panas_daring | PANAS-X item | wagemutig |
| panas_surprised | PANAS-X item | überrascht |
| panas_strong | PANAS-X item | stark |
| panas_scornful | PANAS-X item | voller Verachtung |
| panas_relaxed | PANAS-X item | entspannt |
| panas_irritable | PANAS-X item | reizbar |
| panas_delighted | PANAS-X item | erfreut |
| panas_inspired | PANAS-X item | angeregt |
| panas_fearless | PANAS-X item | furchtlos |
| panas_disgusted_with_self | PANAS-X item | vor mir selbst geekelt |
| panas_sad | PANAS-X item | traurig |
| panas_calm | PANAS-X item | ruhig |
| panas_afraid | PANAS-X item | ängstlich |
| panas_tired | PANAS-X item | müde |
| panas_amazed | PANAS-X item | verblüfft |
| panas_shaky | PANAS-X item | unsicher |
| panas_happy | PANAS-X item | glücklich |
| panas_timid | PANAS-X item | zaghaft |
| panas_alone | PANAS-X item | allein |
| panas_alert | PANAS-X item | hellwach |
| panas_upset | PANAS-X item | verärgert |
| panas_angry | PANAS-X item | wütend |
| panas_bold | PANAS-X item | kühn |
| panas_blue | PANAS-X item | trübsinnig |
| panas_shy | PANAS-X item | schüchtern |
| panas_active | PANAS-X item | aktiv |
| panas_guilty | PANAS-X item | schuldig |
| panas_joyful | PANAS-X item | freudig |
| panas_nervous | PANAS-X item | nervös |
| panas_lonely | PANAS-X item | einsam |
| panas_sleepy | PANAS-X item | schläfrig |
| panas_excited | PANAS-X item | freudig erregt |
| panas_hostile | PANAS-X item | feindselig |
| panas_proud | PANAS-X item | stolz |
| panas_jittery | PANAS-X item | unruhig |
| panas_lively | PANAS-X item | lebhaft |
| panas_ashamed | PANAS-X item | beschämt |
| panas_at_ease | PANAS-X item | gelassen |
| panas_scared | PANAS-X item | verängstigt |
| panas_drowsy | PANAS-X item | dösig |
| panas_angry_at_self | PANAS-X item | über mich selbst verärgert |
| panas_enthusiastic | PANAS-X item | begeistert |
| panas_downhearted | PANAS-X item | niedergeschlagen |
| panas_sheepish | PANAS-X item | verlegen |
| panas_distressed | PANAS-X item | bedrückt |
| panas_blameworthy | PANAS-X item | tadelnswert |
| panas_determined | PANAS-X item | entschlossen |
| panas_frightened | PANAS-X item | furchtsam |
| panas_astonished | PANAS-X item | erstaunt |
| panas_interested | PANAS-X item | interessiert |
| panas_loathing | PANAS-X item | hasserfüllt |
| panas_confident | PANAS-X item | sebstsicher |
| panas_energetic | PANAS-X item | energiegeladen |
| panas_concentrating | PANAS-X item | konzentriert |
| panas_dissatisfied_with_self | PANAS-X item | mit mir selbst unzufrieden |
| hours_of_sleep_usually | "On average, how many hours do you sleep every night?" | "Wie viele Stunden schlafen Sie durchschnittlich pro Nacht?" |
| hours_of_sleep_last_night | "How many hours did you sleep last night?" | "Wie viele Stunden haben Sie letzte Nacht geschlafen?" |
| vigilance | "How well rested do you feel right now? 1 (Extremely tired) - 9 (Perfectly well rested)" | "Wie ausgeruht fühlen Sie sich in diesem Moment? 1 (äußerst müde) - 9 (vollkommen ausgeruht)" |
| quality_of_sleep | "How well did you sleep last night? 1 (I slept terribly) - 9 (I slept very well)" | "Wie gut haben Sie letzte Nacht geschlafen? 1 (sehr schlecht) - 9 (sehr gut)" |
| thirst | "How well hydrated do you feel right now? 1 (Completely dehydrated) - 9 (Perfectly hydrated)" | "Wie durstig sind Sie in diesem Moment? 1 (äußerst durstig) - 9 (überhaupt nicht durstig)" |
| liters_of_water_daily | "On average, how much water (and other liquids) do you drink every day (in litres)?" | "Wie viel Wasser (oder andere Getränke) trinken Sie durchschnittlich pro Tag (in Liter)?" |
| relative_water_intake | "Comparing to other days did you drink more or less water today? 1 (I drank much less than usual) - 5 (I drank the same amount as usual) 9 (I drank much more than usual)" | "Haben Sie heute mehr oder weniger Wasser getrunken verglichen mit anderen Tagen? 1 (sehr viel weniger als gewöhnlich) - 5 (genauso viel wie gewöhnlich) - 9 (sehr viel mehr als gewöhnlich)" |
| caffeine_daily | "On average, how much caffeinated drinks (coffee, cola, club mate etc.) do you drink every day (in litres - one cup = 0.2 l)?" | "Wie viel koffeinhaltige Getränke (Kaffee, Cola, Club Mate, etc.) nehmen Sie durchschnittlich pro Tag zu sich (in Liter: 1 Glas = 0,2 L)?" |
| relative_caffeine_intake | "Comparing to other days did you drink more or less coffee and other caffeinated drinks (coffee, cola, club mate etc.) today? 1 (I drank much less than usual) - 5 (I drank the same amount as usual) 9 (I drank much more than usual)" | "Haben Sie heute im Vergleich zu anderen Tagen mehr oder weniger koffeinhaltige Getränke (Kaffee, Cola, Club Mate, etc.) zu sich genommen? 1 (sehr viel weniger als gewöhnlich) - 5 (genauso viel wie gewöhnlich) - 9 (sehr viel mehr als gewöhnlich)" |
| systolic_blood_pressure_left | Systolic blood pressure measured on the left arm | n/a |
| diastolic_blood_pressure_left | Diastolic blood pressure measured on the left arm | n/a |
| pulse_left | Pulse measured on the left arm | n/a |
| systolic_blood_pressure_right | Systolic blood pressure measured on the right arm | n/a |
| diastolic_blood_pressure_right | Diastolic blood pressure measured on the right arm | n/a |
| pulse_right | Pulse measured on the right arm | n/a |

### CCPT responses

Location: CCPT.csv

File format: plain text, comma separated values

Responses to the CCPT task are available as a CSV file. Each response for each participant and each visit is described by the presented stimuli, response and response time. A description of all items is given in Table 2.

**Table 2.** Description of the columns of the CCPT.csv file.

| Column name | Description |
| --- | --- |
| subject_id | Participant ID |
| session | Which visit the score corresponds to |
| trial_number | One shape presentation is one trial |
| response | Subject response ("space" or "None") |
| response_time | Subject response in milliseconds measured from stimulus onset |
| trial_ISI | Interstimulus interval |
| trial_shape | Shape and colour of the presented stimulus |

### Mini NYC-Q responses

Location: mini_nyc_q.csv

File format: plain text, comma separated values

Responses to the mini NYC-Q questionnaire are available as a CSV file. Each participant responded to every item eight times (four per visit, one after CCPT task, three after resting state scans). A description of all items is given in Table 3.

**Table 3.** Description of the columns of the mini_nyc_q.csv file.

| Column name | Description | German translation (if applicable) |
| --- | --- | --- |
| subject_id | Participant ID | n/a |
| session | Which visit the score corresponds to | n/a |
| timepoint | When the measurement was taken: 1 - after CCPT, 2 - after first whole-brain rs-fMRI, 3 - after second whole-brain rs-fMRI, 4 - after the prefrontal rs-fMRI | n/a |
| positive | "I thought about something positive" | "habe ich an etwas Positives gedacht." |
| negative | "I thought about something negative" | "habe ich an etwas Negatives gedacht." |
| future | "my thoughts involved future events" | "habe ich an zukünftige Ereignisse gedacht." |
| past | "my thoughts involved past events" | "habe ich an vergangene Ereignisse gedacht." |
| myself | "my thoughts involved myself" | "habe ich über mich selbst nachgedacht." |
| people | "my thoughts involved other people" | "habe ich an andere Menschen gedacht." |
| surroundings | "my thoughts involved my surroundings" | "habe ich über meine derzeitige Umgebung nachgedacht." |
| vigilance | "I was fully awake" | "war ich vollkommen wach." |
| images | "my thoughts were in the form of images" | "hatte ich Gedanken in Form von Bildern." |
| words | "my thoughts were in the form of words" | "hatte ich Gedanken in Form von Worten." |
| specific_vague | "my thoughts were more specific than vague" | "waren meine Gedanken eher spezifisch als vage." |
| intrusive | "my thoughts were intrusive" | "waren meine Gedanken aufdringlich/eindringlich." |

### Anatomical scans

Location: niftis/sub<ID>/session_[1-2]/MP2RAGE*.nii.gz

File format: NIfTI, gzip-compressedSequence protocol: acquisition_protocols/MP2RAGE.pdf

MRI data are available in NIFTI formats. All structural scans have been defaced to protect participants identity. The defacing procedure was performed after DICOM to NIFTI conversion and therefore we do not include DICOM files in this dataset. Different reconstruction images are included:

1. INV1 - first inversion time volume
2. INV1_PHS - phase image for the first inversion
3. INV2 - second inversion time volume
4. INV2_PHS - phase image for the second inversion
5. UNI - unified volume (T1 weighted)
6. T1 - quantitative T1 map (note: T1 time estimates above 4000 ms in the CSF are arbitrarily set to 0 ms when calculated on the scanner)

### Fieldmaps

Location: niftis/sub<ID>/session_[1-2]/fieldmap_[1-2]_[magnitude|phase].nii.gz and dicoms/sub<ID>/session_[1-2]/fieldmap_[1-2]_[magnitude|phase].tar.xv

File format: NIfTI, gzip-compressed and DICOM, gzip-compressed directory

Sequence protocol: acquisition_protocols/fieldmap.pdf

fMRI data are available in NIFTI and DICOM formats. The fieldmap_magnitude files consist of two volumes corresponding to the two readouts at different TEs. The fieldmap_phase is the phase difference image.

### Whole-brain rs-fMRI

Location: niftis/sub<ID>/session_[1-2]/rest_full_brain_[1-2].nii.gz and dicoms/sub<ID>/session_[1-2]/rest_full_brain_[1-2].tar.xz

File format: NIfTI, gzip-compressed and DICOM, gzip-compressed directory

Sequence protocol: acquisition_protocols/rest_full_brain.pdf

fMRI data are available in NIFTI and DICOM formats. Even though the NIFTI headers include all of the DICOM meta information thanks to the use of dcmstack converter, some tools (such as AFNI^36^) use different formats for storing meta information in NIFTI files.

### Prefrontal cortex rs-fMRI

Location: niftis/sub<ID</session_[1-2]/rest_prefrontal.nii.gz and dicoms/sub<ID>/session_[1-2]/rest_prefrontal.tar.xz

File format: NIfTI, gzip-compressed and DICOM, gzip-compressed directory

Sequence protocol: acquisition_protocols/rest_prefrontal.pdf

fMRI data are available in NIFTI and DICOM formats. Even though the NIFTI headers include all of the DICOM meta information thanks to the use of dcmstack converter, some tools (such as AFNI^36^) use different formats for storing meta information in NIFTI files.

### Physiological recordings

Location:

sub<ID>/physio/session_[1-2]/physio_trig_resp_card_oxy_[prefrontal|full_brain_1|full_brain_2].txt and sub<ID>/physio/session_[1-2]/physio_original_[prefrontal|full_brain_1|full_brain_2].acq

File format plain text, gzip-compressed or ACQ

Physiological data were down-sampled to 100 Hz and truncated to start with the first MRI trigger pulse and to end one volume acquisition duration after the last trigger pulse. Data are provided in a four-column (MRI trigger, respiratory trace, cardiac trace, and oxygen saturation), space-delimited text file for each resting state scan. Additionally, the original BIOPAC ACQ files recorded at a sampling rate of 5000Hz are provided. They can be read using the bioread package https://github.com/njvack/bioread.

## Technical Validation

### CCPT

To describe the distribution and variance across visits of the CCPT results we have plotted reaction times and percentage of mistakes each participant made (see Figure 3). Reaction times were stable across time and most participants made very few mistakes suggesting that the task was easy for them.

**Figure 3.**
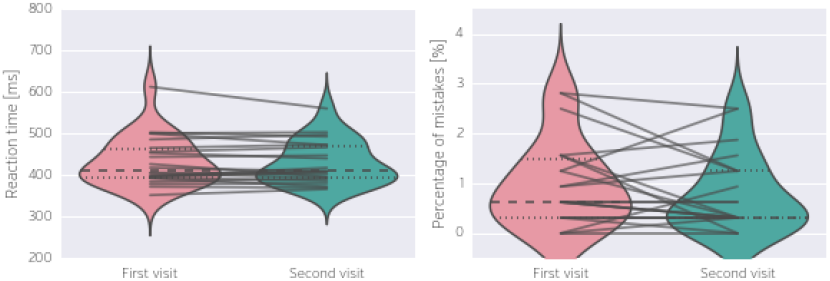
Distribution of reaction times and percentage of mistakes over participants and between two visits. Each line corresponds to one participant. Only correct responses were used to calculate reaction times.

### Mini NYC-Q

To asses the variance of the self reported content of self generated thoughts we have plotted the evolution of the answers over the period of the two visits (see Figure 4). There is a high variance between as well as within participants, consistent with the nature of mind wandering.

**Figure 4.**
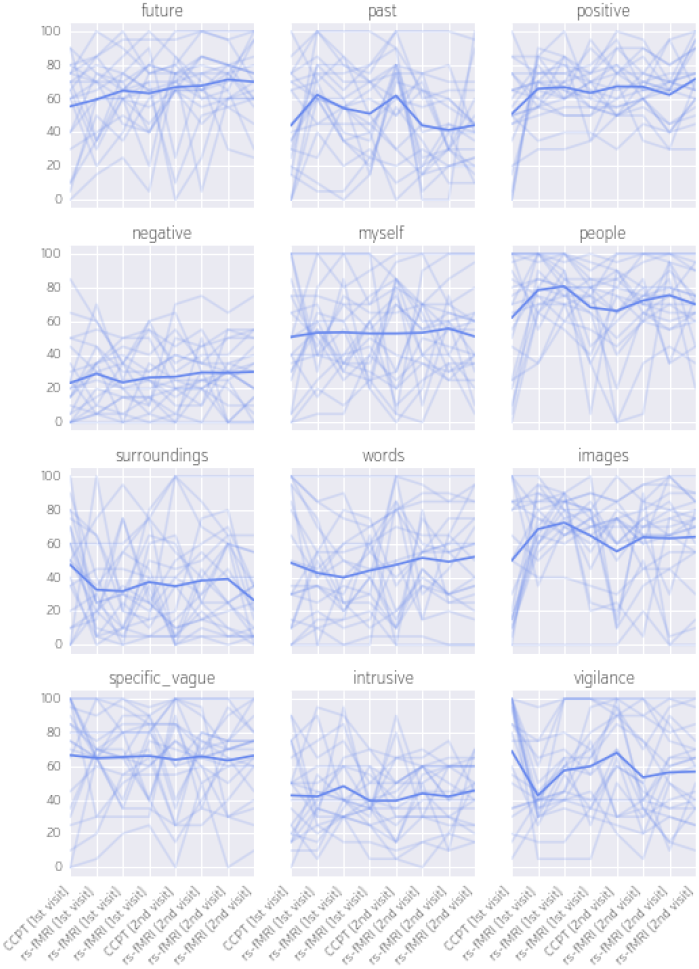
Evolution of the content of self generated thoughts and vigilance over the course of the experiment. Each line corresponds to one participant. First four time points correspond to the first visit - the last four to the second visit. The thick line corresponds to the mean across all participants.

### PANAS-X

We have decomposed the 60 items of the PANAS-X questionnaire into 13 subscales^25^ by averaging corresponding items (see Figure 5). It should be noted that the skewed distributions of “Sadness” and “Hostility” are due to the high responses of the same participant.

**Figure 5.**
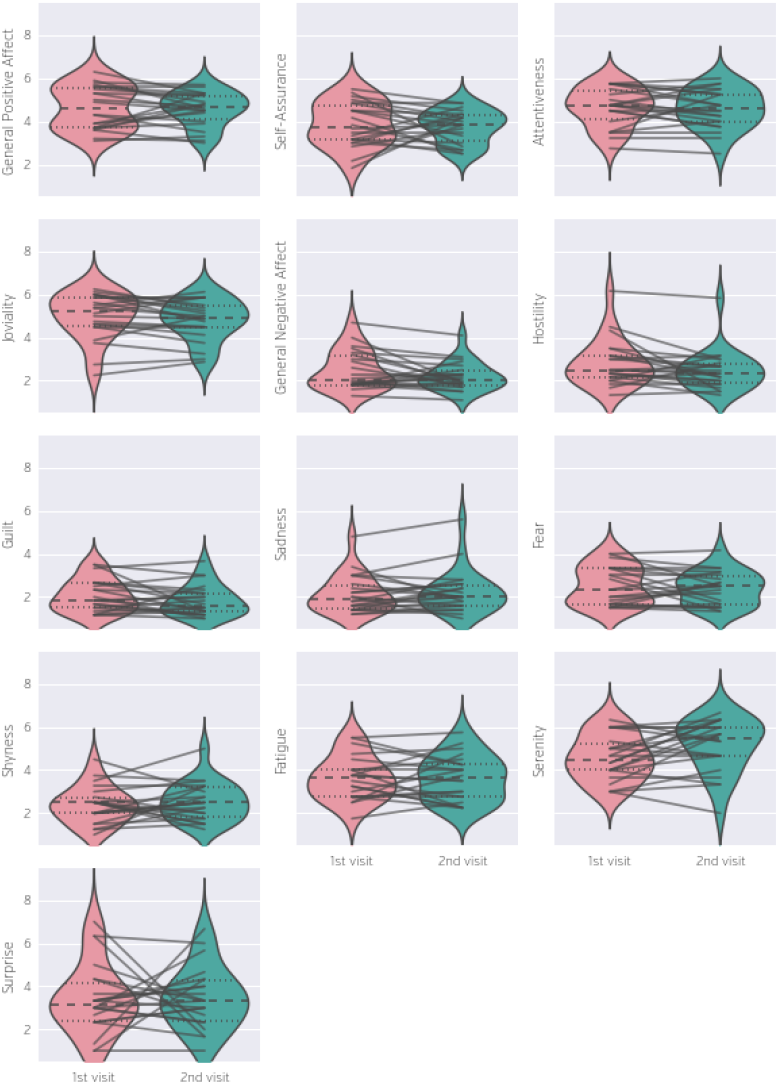
Distribution of PANAS-X subscales scores across participants and between sessions. Score for each subscale was calculated by averaging scores from corresponding items. Each line corresponds to one participant.

### Blood pressure and pulse

As with other measures we also looked at the distribution of blood pressure and pulse measurements across participants and between visits (see Figure 6). Measurements from both arms were averaged together. All participants showed diastolic blood pressure to be within a healthy range (60-79 mm Hg). Some participants, however, experienced systolic blood pressure within the prehypertension range (120-139 mm Hg). The distribution of pulse measurements is rightly skewed due to one outlier (different than the outlier observed in PANAS-X).

**Figure 6.**
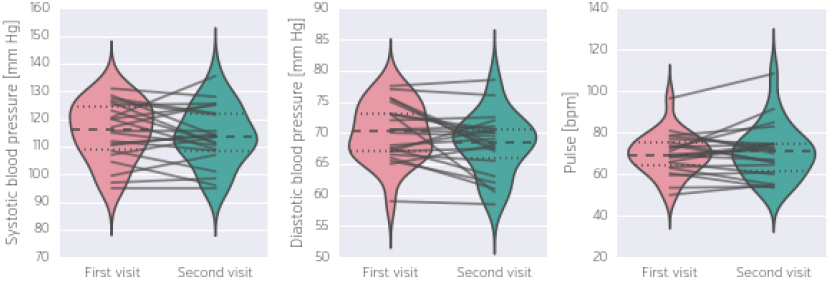
Distribution of systolic and diastolic blood pressure, as well as pulse across participants and between visits. Each line corresponds to one participant.

### Magnetic Resonance Imaging

To asses the quality of the whole-brain resting-state scans we have calculated a series of quality metrics (see Figure 7). Many of these metrics overlap with the ones used by the Consortium for Reliability and Reproducibility (CoRR) (http://fcon_1000.projects.nitrc.org/indi/CoRR) which enables easier comparison with other studies in the consortium. The following metrics were calculated:

1. **Framewise Displacement (FD)** - a measure of volume to volume movement in millimeters. Calculated based on parameters estimated using AFNI *3dvolreg* with *-Fourier -twopass -zpad 4* arguments. Lower values mean less motion.
2. **Temporal Signal to Noise Ratio (tSNR)** - a voxelwise measure of signal strength. It is calculated by dividing the mean across time with standard deviation across time for each voxel individually. A median of all voxels within a brain mask is used to characterise each scan. Mask for this as well as all other metrics was derived using AFNI *3dAutomask* command. Higher tSNR values mean better signal.
3. **Entropy Focus Criterion** - Shannon’s entropy is used to summarize the principal directions distribution^37,38^. Higher values indicate the distribution is more uniform (i.e., less noisy).
4. **Foreground to Background Energy Ratio** - ratio of average absolute value intensities within and outside of the brain mask. Higher values mean more clear signal.
5. **Ghost to Signal Ratio** - A measure of the mean signal in the ‘ghost’ image (signal present outside the brain due to acquisition in a particular phase encoding direction) relative to mean signal within the brain^39^. Lower values indicate fewer ghost artefacts.
6. **Percentage of outliers** - The mean fraction of outlier voxels found in each volume using AFNI *3dTout* command in AFNI. Fewer outliers means better quality.
7. **Median Distance Index** - The mean distance (1 - spearman’s rho) between each time-point’s volume and the median volume using AFNI *3dTqual* command. Smaller values mean more homogeneous timeseries.
8. **Image Smoothness** - Smoothness of the image expressed as a millimeter Full Width Half Maximum. Smoothness was estimated using AFNI *3dFWHMx* command.

Due to a mistake, one of the participant’s whole-brain resting-state scans was acquired using a different sequence. This resulted in a change in the resolution from 1.5 mm to 3.0 mm. We kept the data of this participant in the dataset due to usefulness of all the other measurements, but we do not recommend mixing resolutions in fMRI analysis. This and other data acquisition anomalies are listed in Table 4.

**Figure 7.**
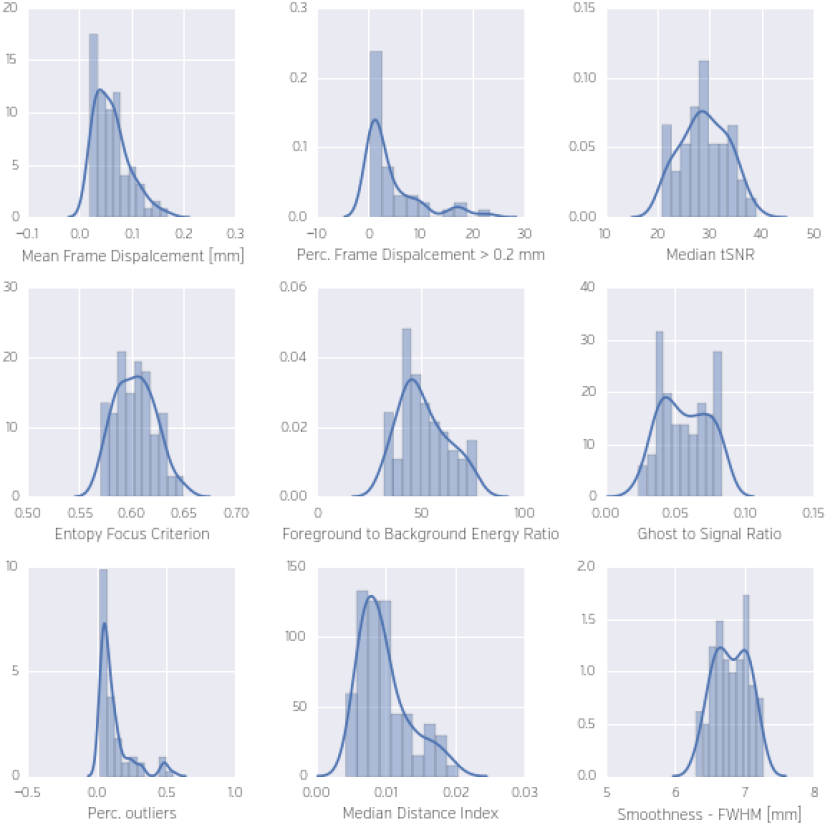
Distribution of quality metrics across whole-brain resting-state scans. One of the participants was excluded due to an acquisition error (wrong scan resolution). Please note that only mean frame displacement and smoothness have meaningful units.

**Table 4.** List of known anomalies in the acquisition process.

| Participant ID | Visit | Issues |
| --- | --- | --- |
| 7 | 1 & 2 | Shimming window was offset causing minor signal deterioration. The same shimming window was used for both sessions. |
| 8 | 1 | The lightbulb in the projector died during the first resting state scan. A projector from another scanner was used as a replacement. The replacement took approximately 30 minutes. The participant was in the scanner during this time. |
| 11 | 1 & 2 | All whole-brain scans were accidentally acquired with voxel size 3mm instead of 1.5mm. |
| 12 | 1 & 2 | Second fieldmap was run after second whole-brain not before, the same order was used for the second session. |
| 13 | 1 | Second magnitude image of the first fieldmap was damaged during transfer from the scanner (one slice is missing). The phase image is intact so the fieldmap can still be reconstructed. |
| 19 | 1 | Second physiological recording (corresponding to second whole-brain scan) was stopped before the scan finished, third physiological recording (corresponding to the prefrontal scan) was started after the scan started. |

## Usage Notes

All data are made available under the terms of the Creative Commons Zero (CC0; PDDL;<u>http://creativecommons.org/publicdomain/zero/1.0/</u>). In short, this means that anybody is free to download and use this dataset for any purpose as well as to produce and re-share derived data artifacts. While not legally enforced, we hope that all users of the data will acknowledge the original authors by citing this publication and follow good scientific practice (i.e. do not try to re-identify the subjects).

Data are shared in documented standard formats, such as DICOM, NIfTI or plain text files, to enable further processing in arbitrary analysis environments with no imposed dependencies on proprietary tools. Many standard software packages may be currently challenged by the increase in data resolution and the changes in anatomical contrasts brought by the MP2RAGE sequence. Dedicated 7T MP2RAGE methods for image segmentation and analysis are freely available within our CBS Tools software package (www.nitrc.org/projects/cbs-tools/) and have been tested and validated on similar data^40^. These tools are modular can be interfaced with other software packages^41^.

## Contributions

KJG conceived and implemented the study and wrote the manuscript. DW, EW, NM, TB, KJG, CG, and DSM acquired the data. RT selected and fine-tuned MRI sequences. FJMR and JS helped with creation of the mini NYC-Q. TB, PLB, DSM, RT, NM and CG helped in writing the manuscript.

